# COVID-19 Severity Is Associated with Differential Antibody Fc-mediated Innate Immune Functions

**DOI:** 10.1101/2021.01.11.426209

**Authors:** Opeyemi S. Adeniji, Leila B. Giron, Netanel F Zilberstein, Maliha W. Shaikh, Robert A Balk, James N Moy, Christopher B. Forsyth, Ali Keshavarzian, Alan Landay, Mohamed Abdel-Mohsen

**Affiliations:** The Wistar Institute, Philadelphia, PA, 19104, USA; Rush University, Chicago, IL, 60612, USA

**Keywords:** SARS-CoV-2, COVID-19, Antibody, Fc-mediated functions

## Abstract

Beyond neutralization, antibodies elicit several innate immune functions including complement deposition (ADCD), phagocytosis (ADCP), and cytotoxicity (ADCC). These functions can be both beneficial (by clearing pathogens) and/or detrimental (by inducing inflammation). We tested the possibility that qualitative differences in SARS-CoV-2 specific antibody-mediated innate immune functions contribute to Coronavirus disease 2019 (COVID-19) severity. We found that antibodies from hospitalized COVID-19 patients elicited higher ADCD but lower ADCP compared to antibodies from non-hospitalized COVID-19 patients. Consistently, higher ADCD was associated with higher systemic inflammation during COVID-19. Our study points to qualitative, differential features of anti-SARS-CoV-2 antibodies as potential contributors to COVID-19 severity.

## INTRODUCTION

While most SARS-CoV-2 infected individuals exhibit asymptomatic or mild respiratory tract infection, a significant population face severe symptoms requiring hospitalization [1, 2]. Hospitalized Coronavirus Disease 2019 (COVID-19) is associated with a state of hyper-inflammation and increased complement activation [1, 3–5]. However, the mechanisms that contribute to this hyper-inflammation are not fully clear.

Higher titers of SARS-CoV-2 specific neutralizing antibodies have been associated with higher COVID-19 severity [6, 7]. However, beyond neutralization, antibodies can elicit an array of Fc-mediated innate immune functions such as antibody-dependent complement deposition (ADCD), antibody-dependent cellular phagocytosis (ADCP), and antibody-dependent cell-mediated cytotoxicity (ADCC) [8]. These innate immune functions can be both beneficial (by clearing pathogens) and/or detrimental (by inducing inflammation) during viral infections [8]. Here, we hypothesized that differential, qualitative features of SARS-CoV2 specific antibodies contribute to COVID-19 severity. To test this hypothesis, we examined the ability of S1- and RBD-specific immunoglobulin G (IgG) from hospitalized and non-hospitalized COVID-19 patients (and controls) to elicit ADCD, ADCP, and ADCC.

## METHODS

### Ethics

All research protocols of the study were approved by the institutional review boards (IRB) at Rush University and The Wistar Institute. All human experimentation was conducted in accordance with the guidelines of the US Department of Health and Human Services and those of the authors’ institutions.

### Characteristics of the study cohort

We isolated IgG antibodies (using Pierce Protein G Spin Plate; Thermo Fisher) from 60 individuals who tested positive for SARS-CoV-2 (by PCR) and 20 negative controls. The 60 SARS-CoV-2 positive individuals were either outpatients (non-hospitalized; n=20) or inpatients (hospitalized; n=40) (**Supplementary Table 1**). Individuals were selected to have a median age between 52.5 to 58.5 years to avoid age bias in disease outcome. The cohort was also chosen to be 45–60% female per group. Samples from hospitalized patients were collected when the patient was admitted. Purified IgG antibodies were diluted with PBS and adjusted to a 1 mg/ml concentration which were used to examine ADCD, ADCP, and ADCC.

### ADCD measurements

Purified IgG antibodies were analyzed for their ability to fix complement on target cells. 4 × 10^6^ (ACE2)-CHO cells were pulsed with 20 μg biotinylated SARS-CoV-2 S1 or RBD proteins (Acro Biosystems) for 30 min at 37°C. Excess, unbound antigens were removed by washing cells once with complete medium. 20 μg IgG was added to the antigen-pulsed cells and incubated for another 30 min at 37°C. Freshly resuspended lyophilized guinea pig complement (Cedarlane) diluted 1:20 with veronal buffer 0.1% gelatin with calcium and magnesium (Boston BioProducts) was added to the cells for 2 h at 37°C. Following a wash with 1X PBS, cells were assessed for complement deposition by staining with goat anti-guinea pig C3-FITC (MP biomedicals). After fixing, cells were analyzed by flow cytometry, and ADCD is reported as MFI of FITC+ cells.

### ADCP measurements

Purified plasma IgG antibodies were analyzed for their ability to mediate phagocytosis of S1 and RBD antigen-coated beads. Biotinylated SARS-CoV-2 S1 and RBD proteins were combined with fluorescent NeutrAvidin beads (Life Technologies) overnight at 4°C. Excess, unconjugated antigens were removed by washing the beads twice with 0.1% PBS-BSA. Beads were washed with 1 ml buffer and spun at 14,000 × g for 2 min at room temperature. Antigen-coated beads were resuspended in a final volume of 1 ml in PBS-BSA. Beads (10 μl) were added into each well of a round-bottom 96-well culture plate, after which 20 μg IgG was added to each well and incubated for 2 h at 37°C. A 200 μl suspension of THP-1 cells at 2.5 × 10^5^ cells/ml was added to each well, for a total of 5 × 10^4^ THP-1 cells per well. After mixing, the cell-bead mixtures were incubated overnight at 37°C. The following day, 100 μl of supernatant from each well were removed, and 100 μl of BD Cytofix were added to each well. Cells were analyzed by flow cytometry, and data collected were analyzed in FlowJo software. The percentage of fluorescent/bead+ cells and the median fluorescence intensity of the phagocytic cells were computed to determine the phagocytosis score.

### ADCC measurements

ADCC induction was estimated by NK cell degranulation and intracellular cytokine production after exposure of antibodies to S1- and RBD-pulsed target cells. 2.5 × 10^4^ S1- or RBD-pulsed (ACE2)-CHO cells were mixed with 20 μg IgG and incubated for 30 min at 37°C. 1 × 10^5^ human NK cells isolated from PBMC using EasySep human NK cell isolation kit (Stem Cell Technologies) were added to wells in the presence of CD107a PE (BD) and Golgi stop (BD). Pulsed (ACE2)-CHO immune complexes were then added to the wells, mixed, pelleted, and incubated at 37°C for 16 h. Post incubation, NK cell activation was detected by staining for CD56 PerCP-Cy5.5 (BD), IFN-γ (BD), TNF (BioLegend) and acquired via flow cytometry. Data are reported as the percentage of cells positive for the marker as indicated.

### Measurement of S1- and RBD-specific antibody titers

S1- and RBD-specific antibody titers were measured in the plasma of the 80 individuals using MSD V-PLEX multiplex assay (Meso Scale Diagnostic).

### Measurement of plasma markers of inflammation

Plasma levels of GM-CSF, IFN-β, IFN-γ, IL-10, IL-13, IL-1β, IL-33, IL-4, IL-6, TNF-α, Fractalkine, IL-12p70, IL-2, IL-21, IL-22, IL-23, IP-10, MCP-2, MIP-1α, SDF-1a, IFN-α2a, IL-12/IL-23p40, and IL-15 were determined using customized MSD U-PLEX multiplex assay (Meso Scale Diagnostic). Plasma levels of C-Reactive Protein (CRP), Galectin-1, Galectin-3, Galectin-9, Growth Differentiation Factor-15 (GDF-15), soluble CD14 (sCD14), soluble CD163 (sCD163), LPS Binding Protein (LBP), and FABP2/I-FABP were measured using ELISA kits (R&D Systems). The plasma level of zonulin was measured using an ELISA kit from MyBiosorce. Levels of occludin were measured by ELISA (Biomatik). Plasma levels of Reg3A were measured by ELISA (RayBiotech). β-glucan detection in plasma was performed using Limulus Amebocyte Lysate (LAL) assay (Glucatell Kit, CapeCod).

### Statistical analysis

Kruskal-Wallis and Mann–Whitney U tests were used for unpaired comparisons. Spearman’s rank correlations were used for bivariate correlation analyses. Statistical analyses were performed in Prism 7.0 (GraphPad).

## RESULTS AND DISCUSSION

### Severe COVID-19 is associated with higher ADCD and lower ADCP compared to mild COVID-19

IgG from hospitalized COVID-19 patients elicited significantly higher ADCD against S1- and RBD-coated target cells compared to IgG from non-hospitalized individuals (**Figure 1a, b**). In contrast, a higher fraction of IgG from non-hospitalized COVID-19 patients elicited ADCP activity above background (maximum value from the SARS-CoV-2 negative group) compared to hospitalized patients (**Figure 1c-e**). S1-specific antibodies from hospitalized individuals induced significantly higher NK cell degranulation and intracellular cytokine production (a surrogate of ADCC) than did S1-specific antibodies from non-hospitalized individuals (**Fig. 1f, g**). In contrast, RBD-specific antibodies from non-hospitalized individuals induced significantly higher ADCC surrogates than did RBD antibodies from hospitalized individuals (**Fig. 1h, i**). These data suggest that hospitalized COVID-19 is associated with differential, qualitative features of SARS-CoV-2-specific antibodies. In particular, hospitalized COVID-19 is associated with a higher ability of antibodies to elicit complement deposition and a lower ability to elicit phagocytosis. These data are compatible with reports suggesting that complement immune system plays a significant negative role in Coronavirus disease pathogenesis [9–11].

**Figure 1.**
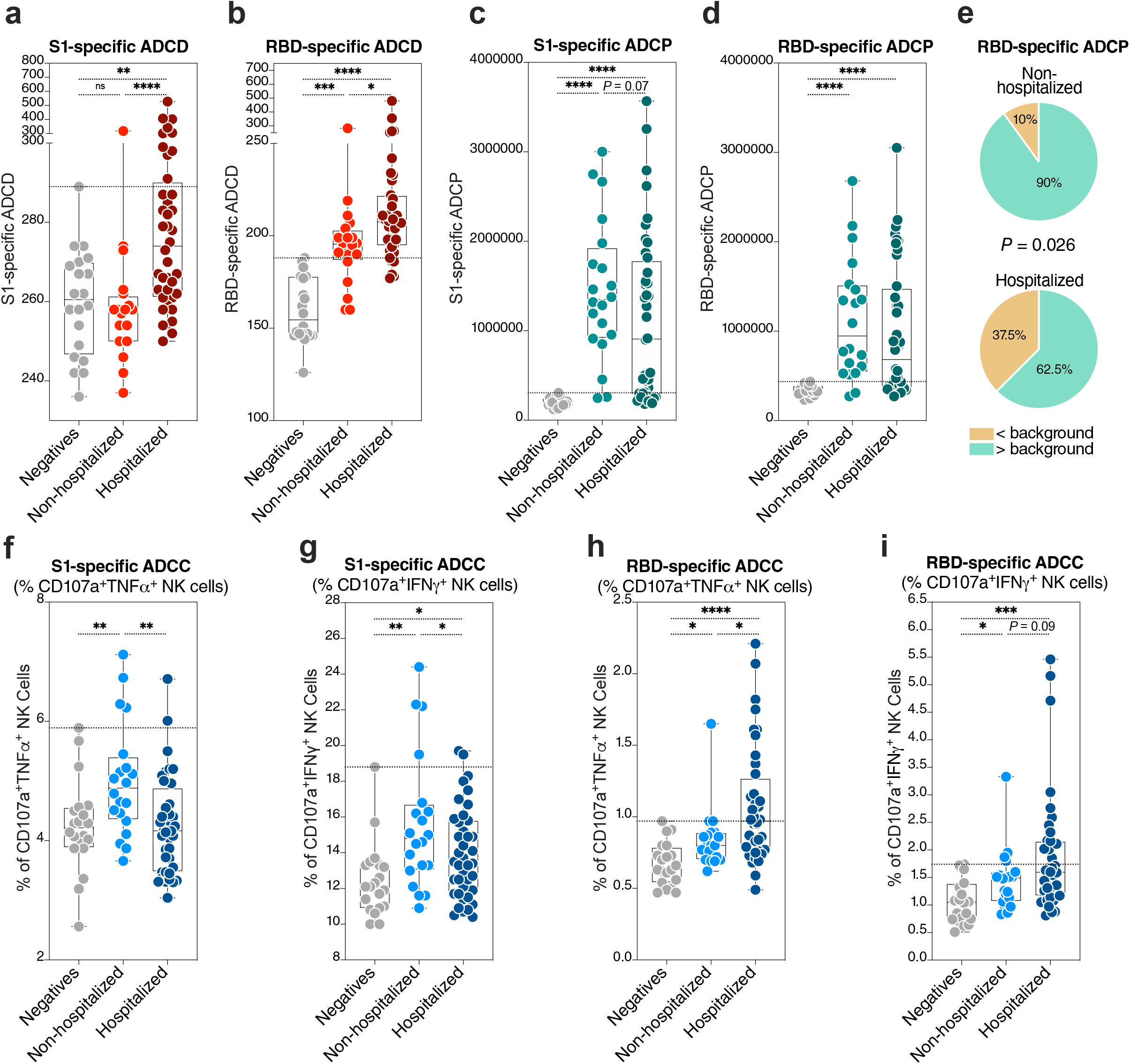
Hospitalized COVID-19 is associated with differential antibody Fc-mediated innate immune functions. **(a-b)** The ability of anti-S1 (a) and anti-RBD (b) antibodies to elicit antibody dependent complement deposition (ADCD) was measured as the ability of 1 mg/ml of IgG to fix complement on target cells. Kruskal–Wallis test was used for statistical analysis. Horizontal dotted lines indicate the maximum values from the SARS-CoV-2 negative control group. **(c-d)** The ability of anti-S1 (a) and anti-RBD (b) antibodies to elicit antibody dependent cellular phagocytosis (ADCP) was measured as the ability of 1 mg/ml of IgG to mediate phagocytosis of S1 and RBD antigen-coated beads. Kruskal–Wallis test was used for statistical analysis. Horizontal dotted lines indicate the maximum values from the SARS-CoV-2 negative control group. **(e)** Comparison between the fraction of non-hospitalized and hospitalized COVID-19 patients whose antibodies elicit anti-RBD-specific ADCP with values higher than background (maximum value from the SARS-CoV-2 negative control group). The Chi-Square test was used for statistical analysis. **(f-i)** The ability of anti-S1 (a) and anti-RBD (b) antibodies to elicit antibody-dependent cell-mediated cytotoxicity (ADCC) was estimated by NK cell degranulation and intracellular cytokine (TNFα or IFNγ) production after exposure of 1 mg/ml of IgG to S1- and RBD-pulsed target cells. Kruskal–Wallis test was used for statistical analysis. Horizontal dotted lines indicate the maximum values from the SARS-CoV-2 negative control group. * = p< 0.05; ** = p<0.01; *** = p<0.001; **** = p<0.0001.

### SARS-CoV-2 antibody titers correspond to ADCP but not ADCD or ADCC

We next sought to examine whether the differential levels of ADCD, ADCP, and ADCC between hospitalized and non-hospitalized patients can be simply explained by the titers of anti-S1 and anti-RBD antibodies. There was no significant difference in the titer of S1-specific antibodies between hospitalized and non-hospitalized patients in our cohort (**Fig. 2a**). When we examined S1-specific antibody titers in patients having ADCD, ADCP, or ADCC values below or above background levels (defined as the maximum values observed in the SARS-CoV-2 negative group), we found no difference in ADCD or ADCC but a significant difference in ADCP (**Fig. 2b-d**). Similar results were observed with RBD-specific antibodies (**Fig. 2e-h**). These data indicate that the quantities of S1- or RBD-specific antibodies could explain the differences in ADCP activity between hospitalized and non-hospitalized patients (the higher the antibody titer, the higher the ADCP activity) but cannot fully explain the differences in ADCD or ADCC activity between hospitalized and non-hospitalized patients, suggesting that qualitative rather than quantitative properties of anti-SARS-CoV-2 antibodies associate with disease severity.

**Figure 2.**
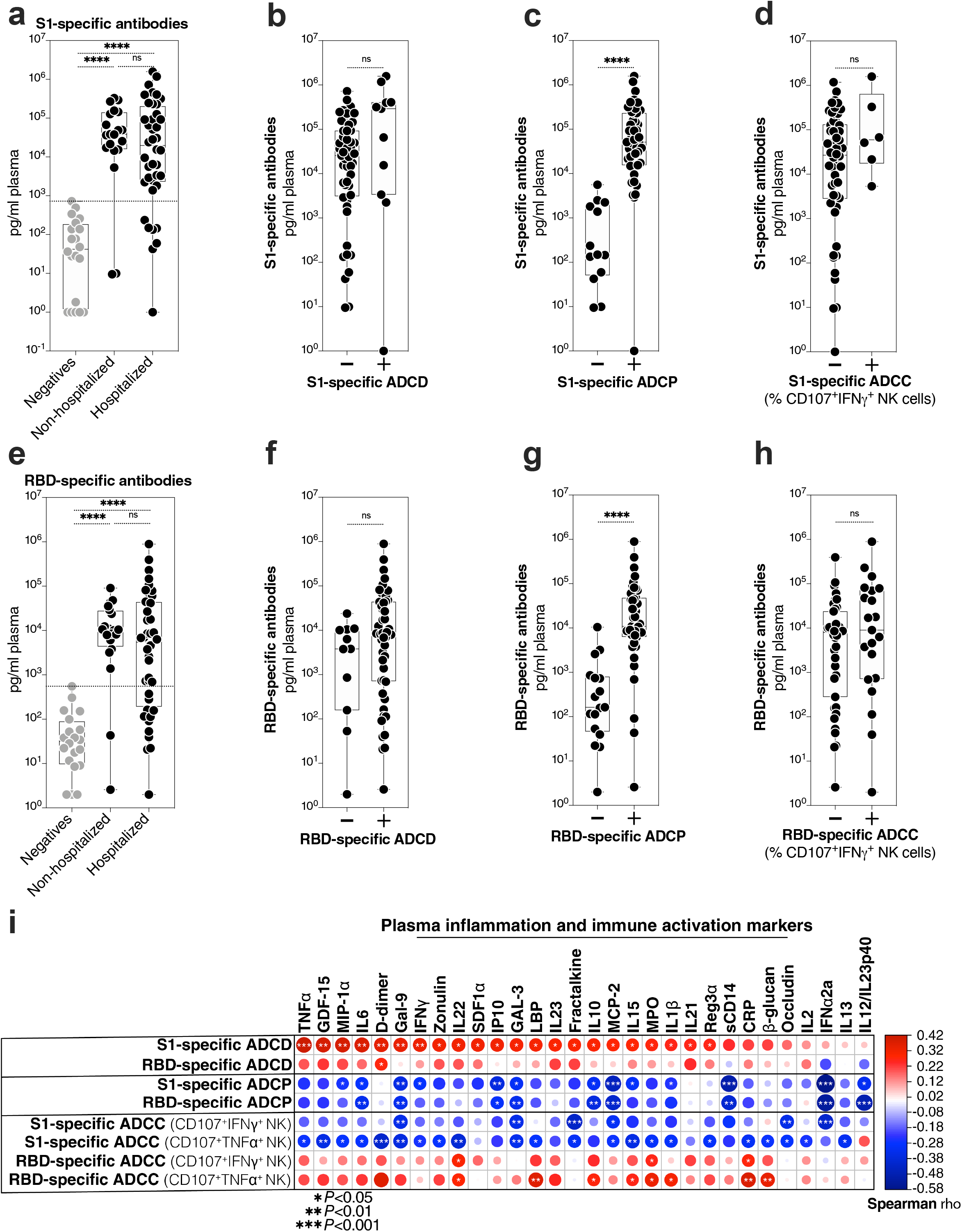
Antibody titers do not correlate with ability to elicit ADCD, which associates with higher systemic inflammation. **(a)** S1-specific antibody titers of the indicated groups. Kruskal–Wallis test was used for statistical analysis. The horizontal dotted line indicates the maximum value from the SARS-CoV-2 negative control group. **(b-d)** S1-specific antibody titers in individuals whose antibodies can elicit S1 specific ADCD (b), ADCP (c), or ADCC (d) higher (+ symbol) or lower (− symbol) than background (maximum value from the SARS-CoV-2 negative control group). Mann–Whitney U test was used in statistical analysis. **(e)** RBD-specific antibodies titers of the indicated groups. Kruskal–Wallis test was used for statistical analysis. The horizontal dotted line indicates the maximum value from the SARS-CoV-2 negative control group. **(f-h)** RBD-specific antibody titers in individuals whose antibodies can elicit RBD-specific ADCD (f), ADCP (g), or ADCC (h) higher (+ symbol) or lower (− symbol) than background (maximum value from the SARS-CoV-2 negative control group). Mann–Whitney U test was used in statistical analysis. **(i)** Correlation heat-map showing associations between Fc-mediated innate immune functions in rows and levels of plasma inflammatory markers in columns during COVID-19 (N=60). The size and color of circles represent the strength of the correlation, with blue shades represent negative correlations and red shades represent positive correlations. * = p< 0.05; ** = p<0.01; *** = p<0.001; **** = p<0.0001.

### Anti-SARS-CoV-2 ADCD associates with higher inflammation

Given that complement activation can significantly contribute to inflammation-mediated tissue damage, including during COVID-19 [12, 13], we examined the relationship between the ADCD property of IgGs (as well as ADCP and ADCC) and an array of 39 plasma markers of systemic inflammation and immune activation measured using ELISA and multiplex cytokine arrays (a list in **Supplementary Table 2**). As expected, higher S1- and RBD-specific ADCD was associated with higher levels of several markers of inflammation and immune activation (**Fig. 2f**). In contrast, higher levels of S1- and RBD-specific ADCP were associated with lower levels of several inflammatory markers (**Fig. 2f**). Finally, ADCC exhibited an antigen-specific relationship with inflammation: anti-S1 specific ADCC exhibited negative correlations with inflammation, whereas anti-RBD exhibited positive correlations with inflammation (**Fig. 2f**). These data further point to the potentially detrimental effects of ADCD on COVID-19 pathogenesis.

Future studies will be needed to examine other antigen-specific antibodies including anti-N antibodies, as well as different antibody classes such as IgA and IgM. These studies will also need to examine the mechanisms that underlie these differential, qualitative features. Antibody glycosylation and subclass have been shown as determinants of Fc-mediated innate immune functions [14]. COVID-19 has been associated with high levels of pro-inflammatory IgG glycomic structures [15]. Understanding how these glycomic features impact the different antibody-mediated innate immune functions during COVID-19 will be needed. Our sample size did not allow for addressing potential confounders. Independent test sets from large cohorts from varying geographic and demographic settings will be needed in future studies. Finally, it will be important to understand how these antibody features impact the function of vaccine-mediated antibodies as well as therapeutic antibodies in order to achieve an optimal balance between neutralization and innate immune functions without inciting potential side effects.

In summary, our data suggest that differential, qualitative Fc-mediated antibody effector properties of anti-SARS-CoV-2 antibodies associate with COVID-19 severity. In particular, antibodies in individuals with hospitalized COVID-19 elicit higher levels of complement deposition compared to antibodies in individuals with mild COVID-19, in a manner linked to higher systemic inflammation. Understanding these qualitative properties will be important to develop COVID-19 therapeutics and SARS-CoV-2 vaccines with optimal efficacy and safety.

---

**Supplementary Table 1.** Demographic and clinical characteristics of the study cohort

**Supplementary Table 2.** A list of plasma markers measured in this study.

## AUTHOR CONTRIBUTIONS

M.A-M conceived and designed the study. O.S.A and L.B.G carried out the experiments. N.F.Z, M.W.S, R.A.B, J.N.M, C.B.F, A.K, and A.L selected study participants and interpreted data. O.S.A and M.A-M wrote the manuscript, and all authors edited it.

## ACKNOWLEDGMENTS

A supplement supports this study to R01 DK123733 (R01 DK123733-01S1) for M.A-M, A.L, and A.K and R24 AA026801-02S1 for A.K. M.A-M is also supported by The Foundation for AIDS Research (amfAR) impact grant # 109840-65-RGR, NIH grants (R01 AG062383, R01NS117458, R21 AI143385, and R21 AI129636), and the Penn Center for AIDS Research (P30 AI 045008). We would like to thank Rachel E. Locke, Ph.D., for providing comments.

