## Supplementary Table 1 for "COVID-19 Severity Is Associated with Differential Antibody Fc-mediated Innate Immune Functions"

**Supplementary Table 1.** Demographic and clinical characteristics of the study cohort

| | | SARS-CoV2 Negative - | SARS-CoV2 Positive outpatients | SARS-CoV2 Positive Inpatients |
|---|---|---|---|---|
| Female, *n* (%) | | 10 (50) | 12 (60) | 18 (45) |
| Age, years, median (IQR) | | 55.5 (15.6) | 52.5 (12.75) | 58.5 (6) |
| Body mass index (BMI) | | - | - | |
| | Normal Weight (<25), % | - | - | 15 |
| | Overweight (25-30), % | - | - | 35 |
| | Obese (30-35), % | - | - | 12.5 |
| | Morbidly Obese (>35), % | - | - | 37.5 |
| Pre-diabetes, % | | - | - | 22.5 |
| Diabetes Mellitus (DM), % | | - | - | 42.5 |
| High blood pressure, % | | - | - | 60 |
| Asthma,% | | - | - | 10 |
| Hydroxychloroquine, % | | - | - | 47.5 |
| Remdesivir, % | | - | - | 15 |
| Tocilizumab, % | | - | - | 17.5 |
| Chronic Steroid Use, % | | - | - | 15 |
| Acute Steroid Use, % | | - | - | 30 |
| Plasma IV nutrition, % | | - | - | 0 |
| Enteral nutrition use, % | | - | - | 100 |
| Antibiotic administration, % | | - | - | 72.5 |
| Ethnicity | | - | - | |
| | African American, *n* (%) | - | - | 16 (40) |
| | Hispanic or Latino | - | - | 15 (37.5) |
| | Caucasian, *n* (%) | - | - | 7 (17.5) |
| | Other, *n* (%) | - | - | 1(2.5) |
| | Unknown, *n* (%) | 20 (100) | 20 (100) | 1 (2.5) |
