## Supplementary Table 2 for "COVID-19 Severity Is Associated with Differential Antibody Fc-mediated Innate Immune Functions"

**Supplementary Table 2.** A list of plasma markers measured in this study.

| Marker | Name | Method of Measurement |
|---|---|---|
| CRP | C-reactive protein | ELISA; R&D Systems |
| IP-10 | C-X-C motif chemokine ligand 10 (CXCL10) | Multiplex meso scale cytokine assay |
| MCP-2 | Chemokine (C-C motif) ligand 8 (CCL8) | Multiplex meso scale cytokine assay |
| Fractalkine | chemokine (C-X3-C motif) ligand 1 (CX3CL1) | Multiplex meso scale cytokine assay |
| C3a | Complement component 3a | ELISA; Thermo Fischer |
| d-dimer | D-dimer | ELISA; Thermo Fischer |
| Gal-1 | Galectin-1 | ELISA; R&D Systems |
| Gal-3 | Galectin-3 | ELISA; R&D Systems |
| Gal-9 | Galectin-9 | ELISA; R&D Systems |
| GM-CSF | Granulocyte-macrophage colony-stimulating factor | Multiplex meso scale cytokine assay |
| GDF-15 | Growth/differentiation factor 15 | ELISA; R&D Systems |
| Zonulin | haptoglobin 2 precursor | ELISA; MyBiosorce |
| IFN-β | Interferon beta | Multiplex meso scale cytokine assay |
| IFN-γ | Interferon gamma | Multiplex meso scale cytokine assay |
| IFN-α2a | interferon α2a | Multiplex meso scale cytokine assay |
| IL-10 | Interleukin 10 | Multiplex meso scale cytokine assay |
| IL-12/IL-23p40 | Interleukin 12 p70 | Multiplex meso scale cytokine assay |
| IL-12p70 | Interleukin 12 p70 | Multiplex meso scale cytokine assay |
| IL-13 | Interleukin 13 | Multiplex meso scale cytokine assay |
| IL-1β | Interleukin 1β | Multiplex meso scale cytokine assay |
| IL-2 | Interleukin 2 | Multiplex meso scale cytokine assay |
| IL-21 | Interleukin 21 | Multiplex meso scale cytokine assay |
| IL-22 | Interleukin 22 | Multiplex meso scale cytokine assay |
| IL-23 | Interleukin 23 | Multiplex meso scale cytokine assay |
| IL-33 | Interleukin 33 | Multiplex meso scale cytokine assay |
| IL-4 | Interleukin 4 | Multiplex meso scale cytokine assay |
| IL-6 | Interleukin 6 | Multiplex meso scale cytokine assay |
| IL-15 | Interleukin-12/interleukin 23 p40 | Multiplex meso scale cytokine assay |
| I-FABP | Intestinal fatty-acid binding protein | ELISA; R&D Systems |
| LBP | Lipopolysaccharide binding protein | ELISA; R&D Systems |
| MIP-1α | Macrophage inflammatory protein alpha | Multiplex meso scale cytokine assay |
| MPO | Neutrophil myeloperoxidase | ELISA; Thermo Fischer |
| OCLN | Occludin | ELISA; Biomatik |
| Reg3A | Regenerating Family Member 3 Alpha | ELISA; RayBiotech |
| sCD14 | Soulble CD14 | ELISA; R&D Systems |
| sCD163 | Soulble CD163 | ELISA; R&D Systems |
| SDF-1a | stromal cell-derived factor 1 (SDF1) or C-X-C motif chemokine 12 (CXCL12) | Multiplex meso scale cytokine assay |
| TNF-α | tumor necrosis factor alpha | Multiplex meso scale cytokine assay |
| β-D-glucan | β-D-glucan | Limulus Amebocyte Lysate (LAL) assay; Glucatell Kit, CapeCod |
